# Cytomegalovirus-induced inactivation of TSC2 disrupts the coupling of fatty acid biosynthesis to glucose availability resulting in a vulnerability to glucose limitation

**DOI:** 10.1101/2023.05.17.541212

**Authors:** Matthew H. Raymonda, Irene Rodríguez-Sánchez, Xenia L. Schafer, Leonid Smorodintsev-Schiller, Isaac S. Harris, Joshua Munger

## Abstract

Human cytomegalovirus (HCMV) modulates cellular metabolism to support productive infection, and the HCMV U_L_38 protein drives many aspects of this HCMV-induced metabolic program. However, it remains to be determined whether virally-induced metabolic alterations might induce novel therapeutic vulnerabilities in virally infected cells. Here, we explore how HCMV infection and the U_L_38 protein modulate cellular metabolism and how these changes alter the response to nutrient limitation. We find that expression of U_L_38, either in the context of HCMV infection or in isolation, sensitizes cells to glucose limitation resulting in cell death. This sensitivity is mediated through U_L_38’s inactivation of the TSC complex subunit 2 (TSC2) protein, a central metabolic regulator that possesses tumor-suppressive properties. Further, expression of U_L_38 or the inactivation of TSC2 results in anabolic rigidity in that the resulting increased levels of fatty acid biosynthesis are insensitive to glucose limitation. This failure to regulate fatty acid biosynthesis in response to glucose availability sensitizes cells to glucose limitation, resulting in cell death unless fatty acid biosynthesis is inhibited. These experiments identify a regulatory circuit between glycolysis and fatty acid biosynthesis that is critical for cell survival upon glucose limitation and highlight a metabolic vulnerability associated with viral infection and the inactivation of normal metabolic regulatory controls.

**Importance:** Viruses modulate host cell metabolism to support the mass production of viral progeny. For Human Cytomegalovirus, we find that the viral U_L_38 protein is critical for driving these pro-viral metabolic changes. However, our results indicate that these changes come at a cost, as U_L_38 induces an anabolic rigidity that leads to a metabolic vulnerability. We find that U_L_38 decouples the link between glucose availability and fatty acid biosynthetic activity. Normal cells respond to glucose limitation by down-regulating fatty acid biosynthesis. Expression of U_L_38 results in the inability to modulate fatty acid biosynthesis in response to glucose limitation, which results in cell death. We find this vulnerability in the context of viral infection, but this linkage between fatty acid biosynthesis, glucose availability, and cell death could have broader implications in other contexts or pathologies that rely on glycolytic remodeling, for example, oncogenesis.

## Introduction

Viruses are obligate parasites that rely on their host cells to supply the metabolic resources necessary for productive replication. Further, viruses have evolved numerous mechanisms to modulate the cellular metabolic environment to support productive infection (Goodwin et al., 2015), although many of the mechanisms involved remain poorly understood. Targeting virally-induced or associated metabolic activities has been an effective anti-viral therapeutic strategy. For example, many therapeutics have been developed that target nucleotide metabolic activities during hepatitis B, HIV, human cytomegalovirus, and human simplex virus infection (Andrei et al., 2008; Cochrane, 2006; Goodrich et al., 1993; Saribey & Tarimci, 2009). More recently, additional metabolic targets have been identified, including lipid-modifying enzymes, whose inhibition can attenuate hepatitis B, Zika, or rhinovirus infections (Grazia Martina et al., 2021). The intersection between viral infection and host-cell metabolism remains a potentially fertile area for anti-viral development.

Human cytomegalovirus is a ubiquitous β-herpesvirus with a double-stranded DNA genome of approximately 235 kB in length. Its genome encodes for over 200 open reading frames which are estimated to produce over 700 viral translation products (Stern-Ginossar et al., 2012). While the majority of HCMV-infected individuals remain asymptomatic, the virus is a substantial cause of morbidity and mortality in immunocompromised individuals such as cancer patients and transplant recipients (Fields et al., 2007). In cancer patients, HCMV has been linked to increased disease progression and worse prognosis (El-Shinawi et al., 2013; Rahbar et al., 2013), while infection during organ transplantation can cause allograft rejection and organ failure (Paya et al., 2004; Stratta et al., 1989). HCMV is also a leading infectious cause of birth defects in the United States and Canada (Bate & Cannon, 2011). These congenital infections have been linked to severe and lifelong complications such as microcephaly, deafness, blindness, motor deficits, and intellectual disabilities (Boppana et al., 2005; Fowler et al., 1999; Kong et al., 2012; Leung et al., 2003; Messinger et al., 2020; Rosenthal et al., 2009; Stagno, Reynolds, Amos, et al., 1977; Stagno, Reynolds, Huang, et al., 1977).

Ganciclovir and its derivative valganciclovir are deoxyguanosine analogs and are the main treatment options for HCMV. However, mutations in U_L_97 and U_L_54 result in resistance, which can emerge in immunosuppressed individuals (Eckle et al., 2000; Kim et al., 2012). Additionally, these compounds suffer from poor bioavailability (Nguyen et al., 2001), and their use is associated with toxicities, including thrombocytopenia and anemia (McGavin & Goa, 2001). Given these limitations, the development of novel therapeutics should remain a priority.

HCMV modulates several metabolic activities to support viral infection, including glycolysis (Landini, 1984), the citric acid cycle (Munger et al., 2006), fatty acid biosynthesis (Munger et al., 2008; Spencer et al., 2011), and pyrimidine biosynthesis (DeVito et al., 2014). Further, many of these activities are important for successful infection (DeVito et al., 2014; McArdle et al., 2012; McArdle et al., 2011; Munger et al., 2008), raising the possibility that they could be targeted for therapeutic intervention.

The HCMV U_L_38 protein is necessary and sufficient to activate glycolysis and amino acid consumption (Rodríguez-Sánchez et al., 2019). U_L_38 binds the tuberous sclerosis complex 2 (TSC2) protein (Moorman et al., 2008; Terhune et al., 2007), which plays a critical role in the regulation of mTORC1 activity by inhibiting the activity of the small GTPase Rheb (Garami et al., 2003; Saucedo et al., 2003; Stocker et al., 2003; Zhang et al., 2003). The phosphorylation of TSC2 by various kinases leads to its dissociation from TSC1, the release of Rheb, and the activation of mTORC1 (Ma et al., 2005). The mTOR pathway is a central regulator of cellular growth, metabolism, and survival, and its dysregulation is linked to a variety of human diseases, including cancer, diabetes, and neurodegenerative disorders (Hall, 2008). When U_L_38 binds TSC2, the tuberous sclerosis complex is inactivated as well, leading to mTORC1 activation and subsequent S6K activation (Moorman et al., 2008). The U_L_38-TSC2 interaction has also been found to be important for U_L_38’s ability to induce its metabolic phenotypes (Rodríguez-Sánchez et al., 2019).

Here, we explore how U_L_38 impacts cellular RNA expression and find that U_L_38 induces the accumulation of many metabolic gene transcripts, including those involved in glycolysis and fatty acid biosynthesis. Consistent with this elevated expression of fatty acid biosynthetic enzymes, we find that U_L_38 strongly activates fatty acid biosynthesis. In control cells, our results indicate that fatty acid biosynthesis is slowed upon limiting glucose availability. Notably, U_L_38-mediated induction of fatty acid biosynthesis is insensitive to lower glucose levels. Further, this U_L_38-induced inability to regulate fatty acid biosynthesis results in cell death upon glucose limitation. This sensitivity depends on U_L_38’s interaction with TSC2, and TSC2 knockout cells display similar sensitivity to glucose withdrawal. Collectively, these experiments identify a metabolic vulnerability associated with viral infection and the inactivation of a critical cell growth regulator that emerges in response to the inability to adapt anabolic pathways to nutrient cues.

## Results

### U_L_38 induces the expression of glycolytic and fatty acid biosynthetic genes

U_L_38 is necessary and sufficient to activate numerous aspects of HCMV-induced metabolic remodeling (Rodríguez-Sánchez et al., 2019), but many questions remain about the mechanisms involved. At early time points during HCMV infection, the U_L_38 protein is localized to the nucleus (Terhune et al., 2007), but little is known regarding its impact on cellular gene expression. To address this issue, we examined cellular RNA abundance in fibroblasts stably expressing U_L_38 relative to control cells (Fig 1A). U_L_38 expression significantly altered cellular RNA levels (Fig 1B) and increased the abundance of many genes associated with central carbon metabolism (shown in turquoise) (Fig 1B; Sup File 1). Ontological pathway analysis of U_L_38 up-regulated genes identified several significantly altered gene pathways (Han et al., 2018; Mi & Thomas, 2009), including glycolysis, pentose phosphate pathway, and cholesterol metabolism (Fig 1C). Glycolysis is strongly induced by U_L_38 expression (Rodríguez-Sánchez et al., 2019), and it was the most enriched pathway observed (Fig 1C). Notably, U_L_38 expression induced the accumulation of mRNA for the majority of enzymatic steps in glycolysis, including key rate-determining enzymes such as hexokinase, phosphofructokinase, and pyruvate kinase (Fig 1D). These results suggest that a major component of U_L_38-mediated metabolic remodeling is the increased expression of glycolytic enzymes.

**Figure 1.**
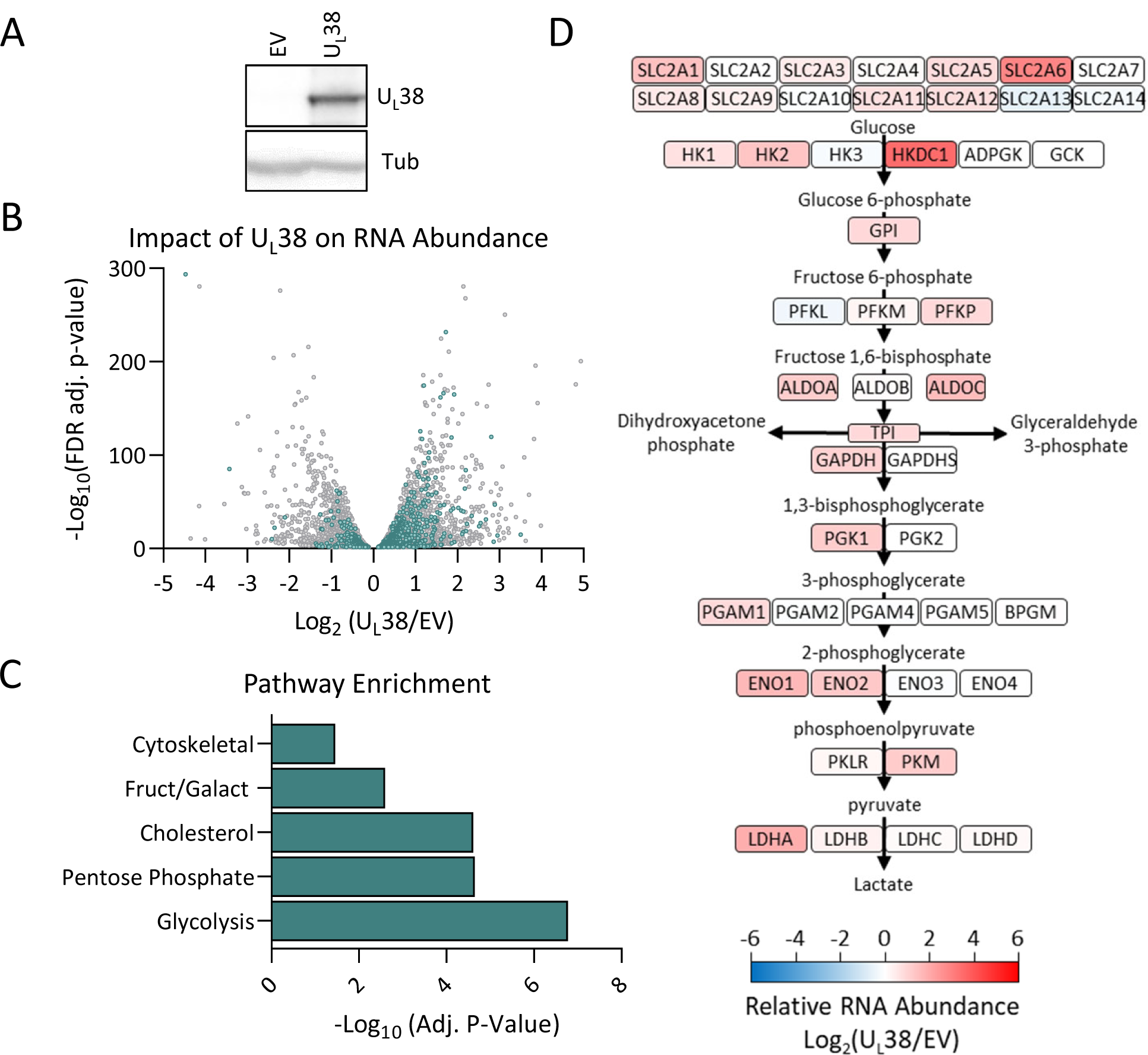
UL38 expression induces the accumulation of central carbon metabolism-associated RNA. Total RNA and protein were harvested from MRC-5 fibroblasts stably expressing either UL38 or an empty vector (EV) and analyzed for protein or RNA abundance. **(A)** Western blot analysis of EV and UL38 expressing cells. **(B)** A volcano plot representing all significant changes in RNA abundance between EV and UL38 expression (padj<0.05). Turquois signifies genes involved in central carbon metabolism. **(C)** Gene set enrichment analysis was performed for all genes with increased abundance in UL38 expressing cells of greater than 50% (Log2(UL38/EV)≥0.6, padj<0.05). **(D)** Diagram of glycolysis pathway from glucose to pyruvate with the Log2(UL38/EV) of individual glycolytic enzymes. Red indicates increased RNA abundance during UL38 expression while blue indicates decreased RNA abundance (n=3).

To identify transcription factors most likely induced by U_L_38 expression, the gene transcripts increased by U_L_38 expression were analyzed via the TRRUST database, which enables the identification of transcriptional relationships between target genes and their transcription factors (Han et al., 2015). This analysis predicted that the sterol regulatory element binding factor 1 (SREBF1) is the most likely U_L_38-activated transcription factor (Fig 2A). U_L_38 expression induced the expression of numerous SREPF1 target genes, including fatty acid synthase (FASN), stearoyl-CoA desaturase (SCD), and the low-density lipoprotein receptor (LDLR) (Fig 2B & S2). SREBF1 has been previously reported to activate lipogenesis during HCMV infection, inducing the expression of multiple proteins involved in sterol synthesis as well as fatty acid biosynthesis (Spencer et al., 2011; Yu et al., 2012). Further, while fatty acid biosynthesis is induced and important for viral replication (Munger et al., 2008; Spencer et al., 2011; Yu et al., 2012), the viral factors responsible have not been identified. Given U_L_38’s induction of fatty acid metabolic enzymes, we tested the hypothesis that U_L_38 is sufficient to activate fatty acid biosynthesis. U_L_38 expression substantially induced fatty acid biosynthesis (Fig 2C). Additionally, we find that U_L_38 is necessary for HCMV-mediated induction of fatty acid biosynthesis (Fig 2D). These results indicate that the U_L_38 protein is a key viral determinant that drives fatty acid biosynthetic activation during HCMV infection.

**Figure 2.**
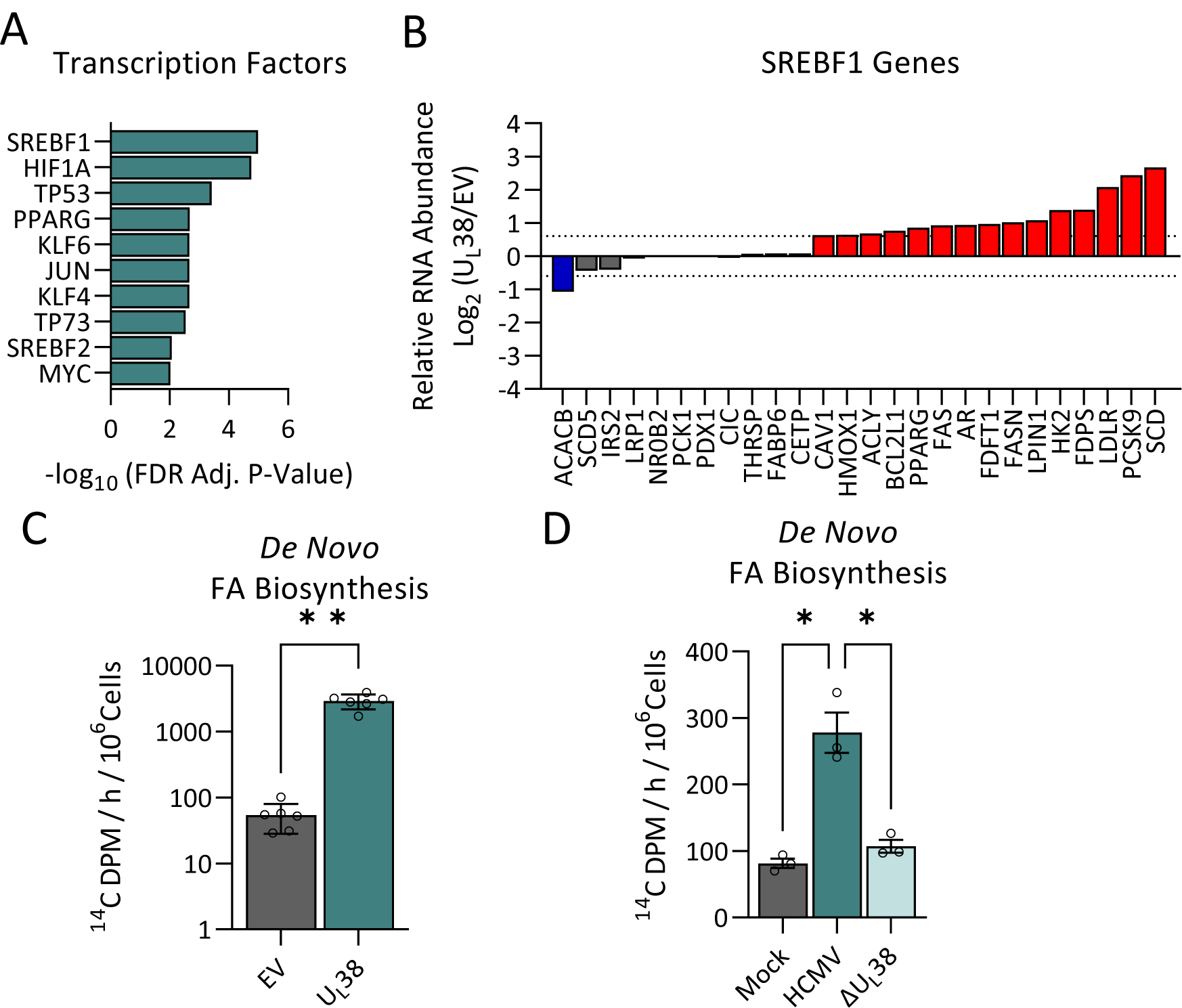
UL38 expression induces fatty acid biosynthesis. **(A)** Transcription factor enrichment analysis from the data in Figure 1B, using RNA abundances that satisfy the following: Log_2_(UL38/EV) ≥ 0.6, (padj<0.05). **(B)** Plot of the relative RNA abundance (Log_2_(UL38/EV)) observed for known gene targets of SREBF1 arranged in order of increasing expression level. Red indicates Log_2_(UL38/EV)≥0.6. Blue indicates Log_2_(UL38/EV)≤-0.6 **(C)** EV or UL38 fibroblasts were treated with 14C-acetate for 4 hours and the rate of fatty acid incorporation measured. Values are means ± SD (n=6). **(D)** MRC-5 cells were mock-infected, infected with WT AD169 (WT), or infected with an AD169 virus lacking UL38 (ΔUL38) at an MOI of 3 PFU/cell. At 48 hours post infection media were replaced with medium containing 14C-acetate for 4h and the rate of fatty acid incorporation measured. Values are means ± SD (n=3).

### Inactivation of TSC2 does not rescue ΔU_L_38 infection

The U_L_38 protein directly interacts with tuberin, a tumor suppressor and negative regulator of the mTORC1 pathway through its role as a GTPase-activating protein for Rheb1. U_L_38 disrupts the tuberous sclerosis complex, which in turn prevents TSC2 from sequestering Rheb1, leading to mTORC1 activation (Moorman et al., 2008). The U_L_38 TQ motif at amino acids 23 and 24 has been shown to promote interaction with TSC2 (Bai et al., 2015; Rodríguez-Sánchez et al., 2019). To determine the importance of the motif upon HCMV infection, we generated an HCMV virus with point mutations (mU_L_38) in this TQ motif, changing it to AA (Bai et al., 2015; Rodríguez-Sánchez et al., 2019). We used this virus to infect fibroblasts and observed that the mU_L_38 protein is expressed at similar levels to WT U_L_38 during infection (Fig 3A). We then examined how the mutant U_L_38 protein impacted glycolysis and fatty acid biosynthesis, both of which are induced by U_L_38 expression alone (Rodríguez-Sánchez et al., 2019) (Fig 2C). We observe that cells infected with mU_L_38 HCMV induce fatty acid biosynthesis and glucose consumption similar to cells infected with WT HCMV (Fig 3B-C). These results suggest that while U_L_38 is sufficient to induce glycolytic activation when expressed in isolation, which depends on its TQ motif, other viral factors are capable of activating glycolysis during HCMV infection when this motif in U_L_38 is mutated. Consistent with a relatively small metabolic phenotype, we do not observe a growth defect in the mU_L_38 virus, in contrast to ΔU_L_38 infection (Fig 3D). These results suggest that the U_L_38’s TQ motif at amino acids 23 and 24 is largely dispensable for high titer replication in fibroblasts.

**Figure 3.**
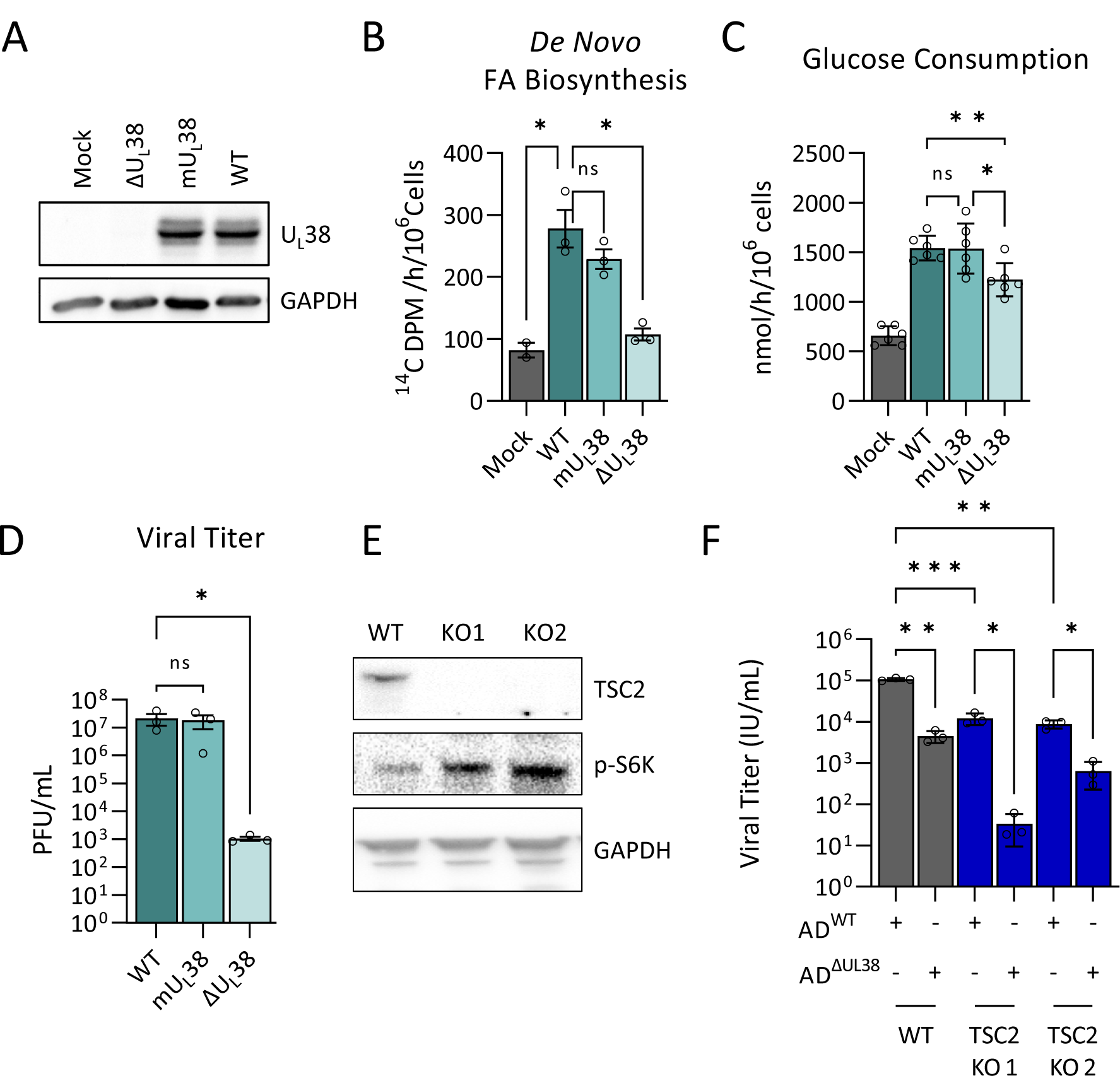
Inactivation of TSC2 does not rescue ΔUL38 HCMV replication. **(A-D)** MRC-5 fibroblasts were mock-infected, infected with WT HCMV, a ΔUL38 mutant, or a mutant expressing the UL38 T23A/Q24A protein variant (mUL38) (MOI = 3). **(A)** At 48 hours post infection, proteins were harvested for western blot analysis. **(B)** At 44 hours post infection media were replaced with medium containing 14C-acetate for 4h. Fatty acids were extracted, saponified and the radioactivity scintillation counted. Values are means ± SD (n=3). **(C)** At 36 hours post infection media were refreshed and harvested 24 hours later and analyzed for glucose abundance. (n=6) **(D)** At 120h post infection, viral media were harvested and titered on MRC-5 via plaque assay. Values are means ± SD (n=3). **(E)** TSC2 knockout MRC5 were generated by electroporating CRISPR Cas9 RNP targeting TSC2 and analyzed by western analysis. **(F)** MRC-5 (WT) and TSC2 KO fibroblasts were infected with WT or ΔUL38 HCMV (MOI = 3). At 120 hours post infection, yields of cell-free virus were titered by IE1 expression on UL38 expressing MRC5 cells. Values are means ± SD (n=3).

To further explore the contributions of TSC2 to viral replication, we generated TSC2 knockout fibroblast cell lines. Delivery of TSC2-directed CAS9 ribonucleoprotein complexes successfully ablated TSC2 protein accumulation (Fig 3E & Fig S2-3). We subsequently assessed whether the inactivation of TSC2 rescues the replication of the ΔU_L_38 mutant. TSC2 ablation did not rescue ΔU_L_38 viral replication (Fig 3G). Further, TSC2 inactivation actually slightly inhibited both WT and ΔU_L_38 infection (Fig 3G). These results suggest that U_L_38’s contribution to infection in fibroblasts is not mediated solely through the inactivation of TSC2.

### U_L_38 expression induces sensitivity to glucose limitation

Glycolytic inhibition attenuates HCMV infection (Landini, 1984; McArdle et al., 2011). In line with these findings, we found that infection in the absence of glucose blocks the production of infectious virions (Fig 4A). This loss of viral production correlated with a reduced abundance of viable cells during infection in the absence of glucose, which did not occur with mock infection (Fig 4B & 4C). Given U_L_38’s role in glycolytic activation, we examined whether the presence of U_L_38 impacted the sensitivity to glucose withdrawal. In contrast to cells infected with WT HCMV, cells infected with the ΔU_L_38 mutant or the mU_L_38 mutant, failed to induce sensitivity to glucose limitation (Fig 4C). These results suggest that U_L_38 expression and its interaction with TSC2 are important for HCMV’s sensitivity to glucose limitation. We subsequently tested whether U_L_38 expression in isolation can induce sensitivity to glucose withdrawal, and found that U_L_38 expression was sufficient to induce sensitivity to glucose limitation (Fig 4D). In contrast, cells stably expressing the mU_L_38 protein did not exhibit sensitivity to glucose limitation (Fig 4E). Collectively, these results suggest that U_L_38 is necessary and sufficient to induce sensitivity to glucose withdrawal and that this sensitivity is modulated based on its ability to attenuate TSC2 function.

**Figure 4.**
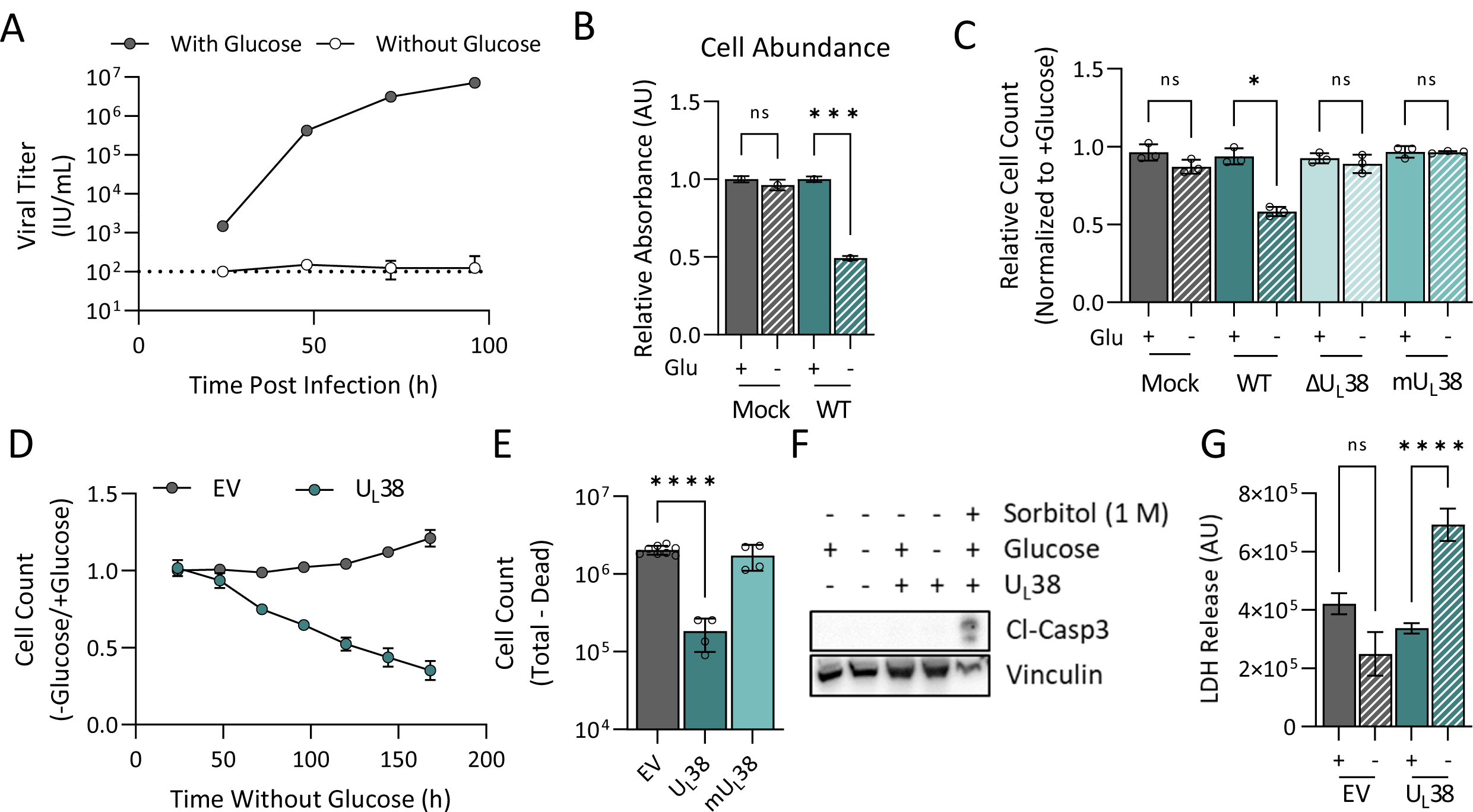
UL38 expression induces sensitivity to glucose limitation. **(A-C)** MRC-5 fibroblasts were mock-infected, or infected with WT HCMV, ΔUL38, or the mUL38 mutant (MOI=3.0). After 2h absorption, media were changed to glucose-containing (+glucose) or glucose-free (-glucose) media. **(A)** Virus containing media were harvested at 24, 48, 72, or 96 hours post infection and yields of cell-free virus were titered by IE1 expression on UL38 expressing MRC5 cells. Values are means ± SD (n=4). **(B)** At 48h post glucose-starvation, media were aspirated, and cell abundance was quantified by crystal violet staining. Absorbances were measured at 595nm and normalized to the absorbance of each sample +glucose control. Values are means ± SD (n=3). **(C)** At 48h post glucose-starvation cellular nuclei were stained, imaged, and counted. The nuclei counts were normalized by the glucose treated control. Values are means ± SD (n=3). **(D)** MRC5 cells expressing an empty vector control (EV) or UL38 protein were grown to confluence and challenged with glucose free medium. At the indicated times, cells were fixed, nuclei stained, and counted. Values are mean ± SD (n=120). **(E)** MRC5 cells expressing an empty vector control (EV), UL38, or mUL38 were grown to confluence and challenged with glucose free medium. At 48h post media change, cells were counted using trypan blue exclusion, dead cells were subtracted from total. Values are mean ± SD (n=4). **(F)** Empty vector and UL38-expressing cells were grown to confluence and challenged with glucose free medium. At 48h post media change, proteins were harvested for western blot analysis. As a positive control for apoptosis, UL38 cells were treated with 1M sorbitol for 1h prior to protein harvesting. Cl-Casp3 = cleaved caspases 3. **(G)** Empty vector and UL38-expressing cells were grown to confluence and challenged with glucose free medium. At 48h post media change, aliquots of media were harvested and analyzed for LDH activity. Values are mean ± SEM (n=15).

It has previously been reported that U_L_38 expression inhibits intrinsic apoptosis (Terhune et al., 2007) and suppresses ER stress-induced cell death (Qian et al., 2011; Xuan et al., 2009). To explore whether the U_L_38-mediated sensitivity to glucose withdrawal involved activation of caspases, we assayed for the accumulation of cleaved caspase 3. In contrast to osmotically shocked cells (induced by sorbitol treatment), the absence of glucose did not increase the levels of cleaved caspase 3 in either EV or U_L_38-expressing cells (Fig 4E). To examine whether the loss of cells was necrotic, we measured LDH release and observed that U_L_38-expressing cells released substantially higher levels of LDH into the media in the absence of glucose, indicative of a loss of membrane integrity and the induction of necrosis (Fig 4G).

### U_L_38 expression alters the metabolic response to glucose limitation

Given the impact of U_L_38 expression on cellular metabolism, we hypothesized that U_L_38 expression would alter the normal response to glucose withdrawal, potentially providing clues about the mechanisms involved. Toward this end, we employed LC-MS/MS to examine the impact of glucose limitation on metabolite pool sizes. Principal component analysis of the metabolite pools indicated that U_L_38 expression substantially alters cellular metabolism relative to empty vector (EV) control cells as shown by sample segregation along both PC1 and PC2 (Fig 5A & S4, Sup File 2-5). Glucose starvation in control cells resulted in a large shift in PC1 (Fig 5A), whereas U_L_38-expressing cells shifted to a much lesser extent along PC1 upon glucose starvation (Fig 5A), indicating that they had a significantly different response to glucose limitation. As would be expected, both control and U_L_38-expressing cells exhibited a reduction in many glycolytic pools, including fructose phosphate, fructose bisphosphate, dihydroxyacetone phosphate (DHAP), and lactate (Fig 5B & 5C, Fig S4). Further, cellular lactate secretion fell in both cell lines upon glucose removal (Fig S6). Glucose-6-phosphate pools were significantly elevated in EV control cells but did not change in U_L_38-expressing cells upon glucose limitation (Fig 5B & 5C). This induction of glucose-6-phosphate pools could reflect increased glycogen breakdown in control cells relative to U_L_38-expressing in response to low glucose levels.

**Figure 5.**
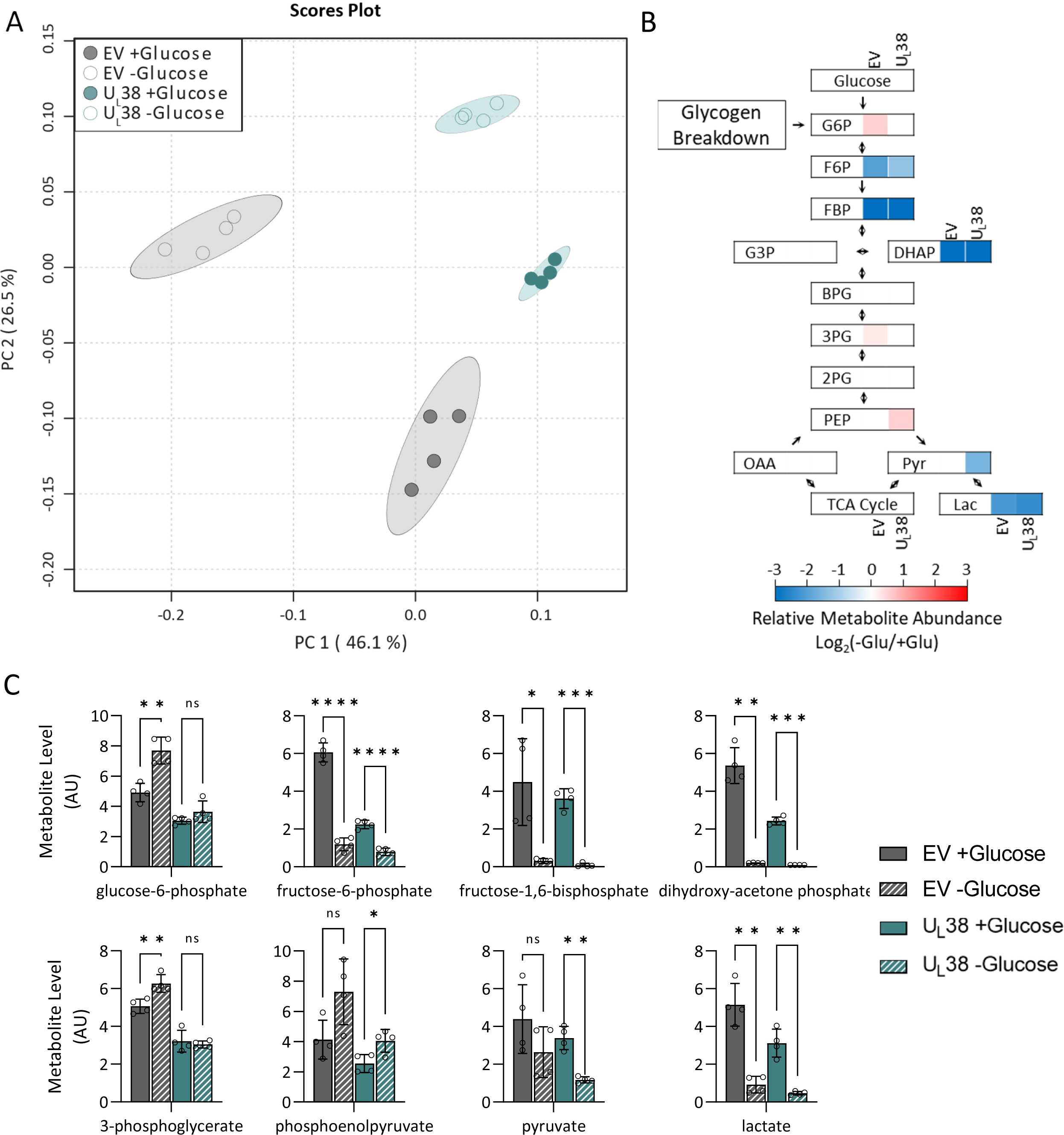
UL38 expression alters the metabolic response to glucose limitation including the abundance of gluconeogenesis-associated metabolites. MRC-5 fibroblasts expressing an empty vector (EV) or wild-type UL38 (UL38) were grown to confluence and challenged with glucose free medium. Media were replaced with glucose-containing or glucose free media for 24 hours and cellular metabolites were harvested and analyzed via LC-MS/MS. **(A)** Principal component analysis showing separation of EV and UL38 with and without glucose. **(B)** Canonical glycolysis pathway displaying changes in metabolite abundance upon glucose withdrawal. If the difference was not significant, the color is white. **(C)** Relative metabolite levels for individual components of glycolysis. Grey = EV, Turquois = UL38, Striped = no glucose. Values are mean ± SD (n=4).

Pyruvate levels trended downward in both cell types but were only significantly reduced in U_L_38-expression cells upon glucose limitation (Fig 5B-5C). The levels of acetyl-CoA, the metabolite immediately downstream of pyruvate, were differentially impacted by the expression of U_L_38. Glucose limitation substantially reduced acetyl-CoA levels in control cells, yet they were increased two-fold in U_L_38-expressing cells (Fig 6A & 6B). Acetyl-CoA supplies two carbon units to the TCA cycle upon citrate synthesis, but acetyl-CoA is also utilized for fatty acid biosynthesis with a major regulatory step being its carboxylation to form malonyl-CoA, the levels of which were induced by U_L_38-expression upon glucose starvation, and which does not occur in control cells (Fig 6A & 6B). These results suggest that U_L_38 expression impacts the metabolic utilization of acetyl-CoA upon glucose limitation.

**Figure 6.**
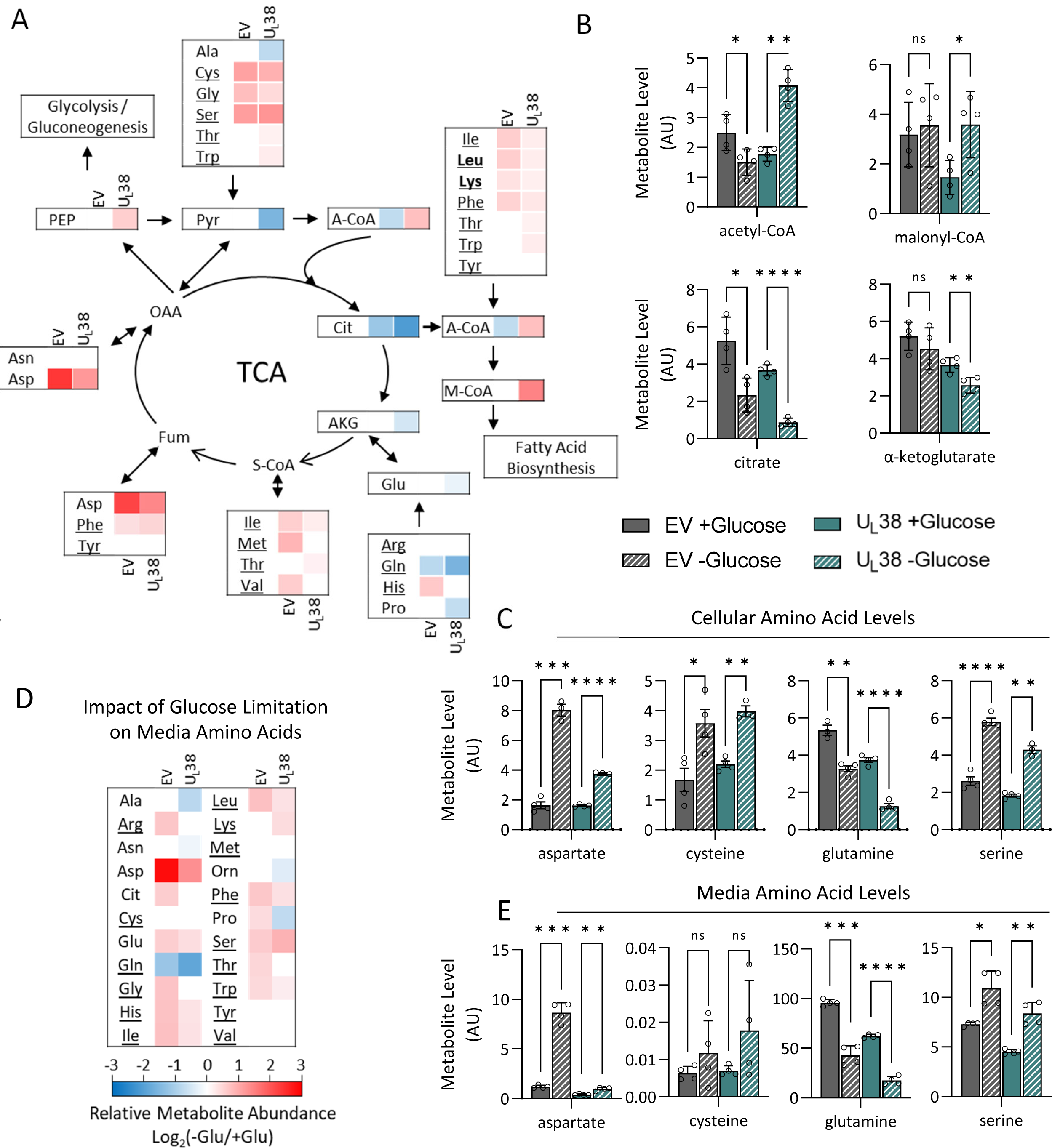
U_L_38 expression induces the abundance of fatty acid biosynthetic precursors during glucose limitation. MRC-5 fibroblasts expressing an empty vector (EV) or wild-type UL38 (UL38) were grown to confluence and challenged with glucose free medium. Media were replaced with glucose-containing or glucose free media for 24 hours and cells and media were harvested and analyzed via LC-MS/MS. **(A)** Diagram of where amino acids feed into the tricarboxylic acid cycle and their relative change in cellular abundance upon glucose withdrawal. If the change was not significant, the color is white. Underlined amino acids are present in the growth media. Bold amino acids are ketogenic only. **(B-C)** Relative intracellular metabolite levels in EV and UL38 cells **(D)** Heatmap of the relative abundance of metabolites in the media 24 hours after media change. **(E)** Relative metabolite levels in the media of EV and UL38 cells. **(B,C,E)** Grey = EV, Turquois = UL38, Striped = no glucose. Values are mean ± SD (n=4).

Glucose limitation reduced the levels of citrate and α-ketoglutarate in both control and U_L_38-expressing cells (Fig 6A & 6B). Further, the levels of cellular glutamine, a major TCA carbon source, were also reduced upon glucose starvation in both conditions (Fig 6A & 6C). Despite this reduction in glutamine levels, both control and U_L_38-expressing cells significantly increased their consumption of media glutamine in response to glucose limitation (Fig 6D & 6E). The induction of glutamine consumption was notable as none of the other amino acids present in the media were consumed at increased rates in either cell line upon glucose limitation (Fig 6D). Glutamine is capable of providing gluconeogenic carbon via TCA-mediated conversion to oxaloacetate and subsequent decarboxylation and phosphorylation to phosphoenolpyruvate via phosphoenolpyruvate carboxykinase (PEPCK) activity. This enhanced gluconeogenic utilization of glutamine could explain the observed maintenance/increases in PEP levels upon glucose limitation (Fig 5C). Gluconeogenic utilization of glutamine requires deamidation of glutamine and deamination of glutamate, both of which result in the production of ammonia, whose accumulation can be toxic. Notably, the levels of aspartate substantially increased upon glucose limitation, especially in the control cells (Fig 6C). Aspartate can potentially serve as an ammonia sink as its synthesis involves a transamination reaction of fumarate or oxaloacetate (Fig 6A). Consistent with this possibility, upon glucose limitation cells increased their export of aspartate, which is absent in fresh media (Fig 6D & 6E). The cellular levels of other amino acids that can feed the citric acid cycle and support gluconeogenesis were also increased by glucose limitation regardless of whether U_L_38 was expressed, including cysteine and serine, (Fig 6A & 6C). For several other amino acids, glucose limitation significantly induced their levels in control cells but did not substantially impact their levels in U_L_38-expressing cells, including histidine, methionine, and valine (Fig S5), indicating that U_L_38 significantly altered the response of these pools to glucose withdrawal.

### Fatty acid biosynthetic inhibitors attenuate U_L_38-mediated sensitivity to glucose limitation

U_L_38 expression induced substantial metabolic changes in response to glucose limitation. To functionally interrogate metabolic pathways involved in U_L_38-induced sensitivity to glucose limitation, we utilized a compound library that focuses on metabolic targets, and which has been previously utilized to examine cancer cell metabolic sensitivities (Harris et al., 2019; Nicholson et al., 2019; Raymonda et al., 2022; Shu et al., 2020). We screened for compounds capable of rescuing cell death associated with U_L_38-mediated sensitivity to glucose limitation (Fig. 7A). Specifically, we used the increase in cell number as an assay of a compound’s ability to inhibit glucose sensitivity. Cell numbers were quantified using an image-based readout and conditions were optimized to generate a robust Z’ factor for the screen (Z’ = 0.49) (Zhang et al., 1999). Each drug in the compound library was arrayed across a 10-point dose curve, ranging from 20 µM to 1 nM, and consisted of multiple compounds targeting each metabolic pathway. A dose-response curve was plotted for each compound and the area under the curve (AUC) of cell number plotted against compound concentration was used to determine effectiveness (Fig. 7A). Of the 240 compounds tested, 60 compounds increased the cellular AUC upon glucose withdrawal. These compounds targeted multiple different pathways including lipogenesis and redox homeostasis (Fig. 7B). Several compounds that targeted components of the fatty biosynthesis pathway were observed to increase cell numbers during glucose withdrawal (Fig. 7C & 7D). These compounds targeted various steps of fatty acid biosynthesis including acetyl-CoA carboxylase (ACC) and fatty acid synthetase (FASN) (Fig. 7E). Three of the hit compounds, ND-630, Fatostatin A, and GSK2194069 were further analyzed for cell death inhibition by analyzing their impact on LDH release over time upon glucose removal. ND-630 and Fatostatin A reduced LDH release at every dose tested (Fig 7F). GSK2194069 was cytotoxic at 25 µM but prevented LDH release in response to glucose limitation at 2.5 and 0.25 µM (Fig 7E). These results indicate that fatty acid biosynthesis is necessary for U_L_38-induced sensitivity to glucose limitation.

**Figure 7.**
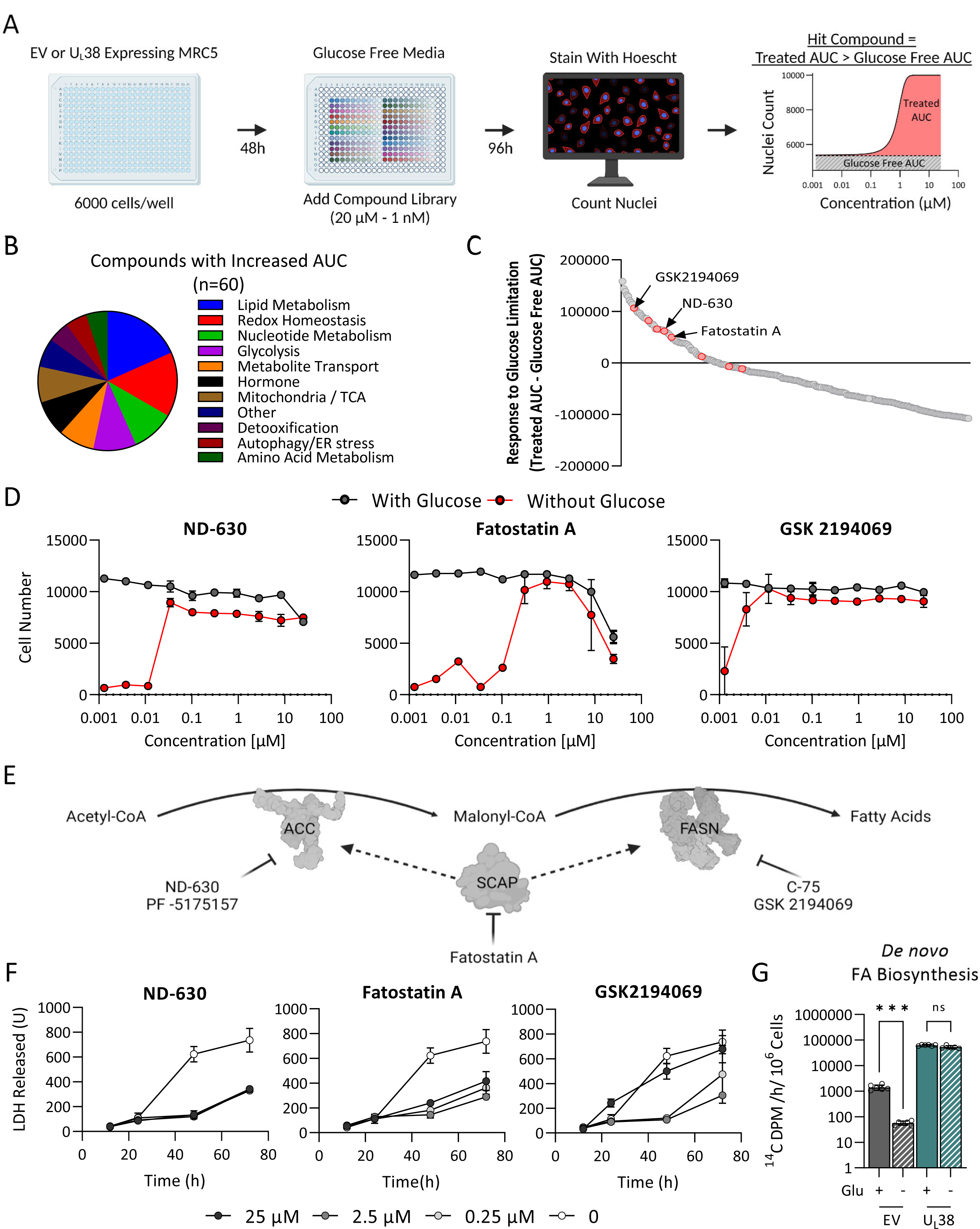
Inhibition of fatty acid biosynthesis attenuates UL38-mediated sensitivity to glucose withdrawal. **(A)** The screening platform used to detect compounds that inhibit glucose sensitivity. Fibroblasts grown in 384-well plates were challenged with glucose free medium and then treated with our compound library. The ability of the compounds to inhibit glucose sensitivity was measured 96 hours after glucose withdrawal by nuclei counting. Hit compounds were determined by comparing the AUC of the nuclei counts to untreated glucose-free counts. **(B)** Pathways associated with the 60 compounds observed to inhibit glucose sensitivity compared to glucose-free control. **(C)** Ranking of library compounds according to the AUC of nuclei counts under glucose-free conditions. The AUC of UL38 cells in the absence of glucose is equal to 0. Red dots represent compounds targeting fatty acid biosynthesis **(D)** Screen data of select fatty acid biosynthesis inhibitors ND-630, Fatostatin A, and GSK-2194069. Grey curves indicate cells treated with compounds in glucose containing media. Red curves indicate cells treated with the compounds in glucose-free media. (n=2) **(E)** Schematic of fatty acid biosynthesis showing where select inhibitors interact. ACC = Acetyl CoA Carboxylase, SCAP = SREBP Cleavage activating protein, FASN, fatty acid synthase. **(F)** UL38 expressing fibroblasts were given glucose-containing or glucose free media containing increasing concentrations of inhibitors (0, 0.25, 2.5, and 25 µM) and LDH activity was analyzed at 12, 24, 48, and 72 hours. Values are mean ± SD (n=3). **(G)** Empty vector (EV) and wild-type UL38 (UL38) expressing fibroblasts were labeled with 14C-acetate in either glucose or glucose-free media. Lipids were extracted, saponified and radioactivity in the lipogenic fraction scintillation counted. Values are means ± SD (n=6).

Given that fatty acid biosynthetic inhibitors attenuated U_L_38-induced sensitivity to glucose limitation, we hypothesized that U_L_38-expression might prevent the modulation of fatty acid biosynthesis in response to glucose limitation. To explore this possibility, we examined how U_L_38 expression impacted fatty acid biosynthesis in response to glucose limitation. In contrast to control cells, which reduced their fatty acid biosynthetic rates in response to glucose withdrawal, U_L_38-expressing cells maintained fatty acid biosynthesis at rates that were 100-fold higher even in the absence of glucose (Fig 7F). Collectively, these results indicate that U_L_38 expression prevents the normal regulation of fatty acid biosynthesis in response to glucose limitation, thereby resulting in an increased sensitivity to this metabolic challenge.

### TSC2 inactivation induces sensitivity to glucose limitation and decouples fatty acid biosynthesis from glucose availability

Our results suggest that U_L_38-induced sensitivity to glucose limitation is a result of the inability to reduce fatty acid biosynthesis upon glucose withdrawal. Given that the expression of the mU_L_38 allele, which exhibits reduced TSC2 inhibition, does not induce sensitivity to glucose limitation (Fig 4E), our model predicts that cells expressing this mU_L_38 allele will be able to down-regulate fatty acid biosynthesis upon glucose withdrawal. Consistent with this prediction, cells expressing the mU_L_38 allele reduced fatty acid biosynthesis in response to glucose limitation, similar to control cells, and in contrast to cells expressing wtU_L_38 (Fig 8A). These results suggest that TSC2 inactivation is responsible for the inability to regulate fatty acid biosynthesis and cell death upon glucose withdrawal. Consistent with this hypothesis, TSC2 inactivation also resulted in fatty acid biosynthetic insensitivity and cell death in response to glucose limitation (Fig 8B & 8C). These results indicate that TSC2 is necessary for the proper coupling of glucose availability to fatty acid biosynthesis, which is necessary to maintain cellular viability upon glucose limitation.

**Figure 8.**
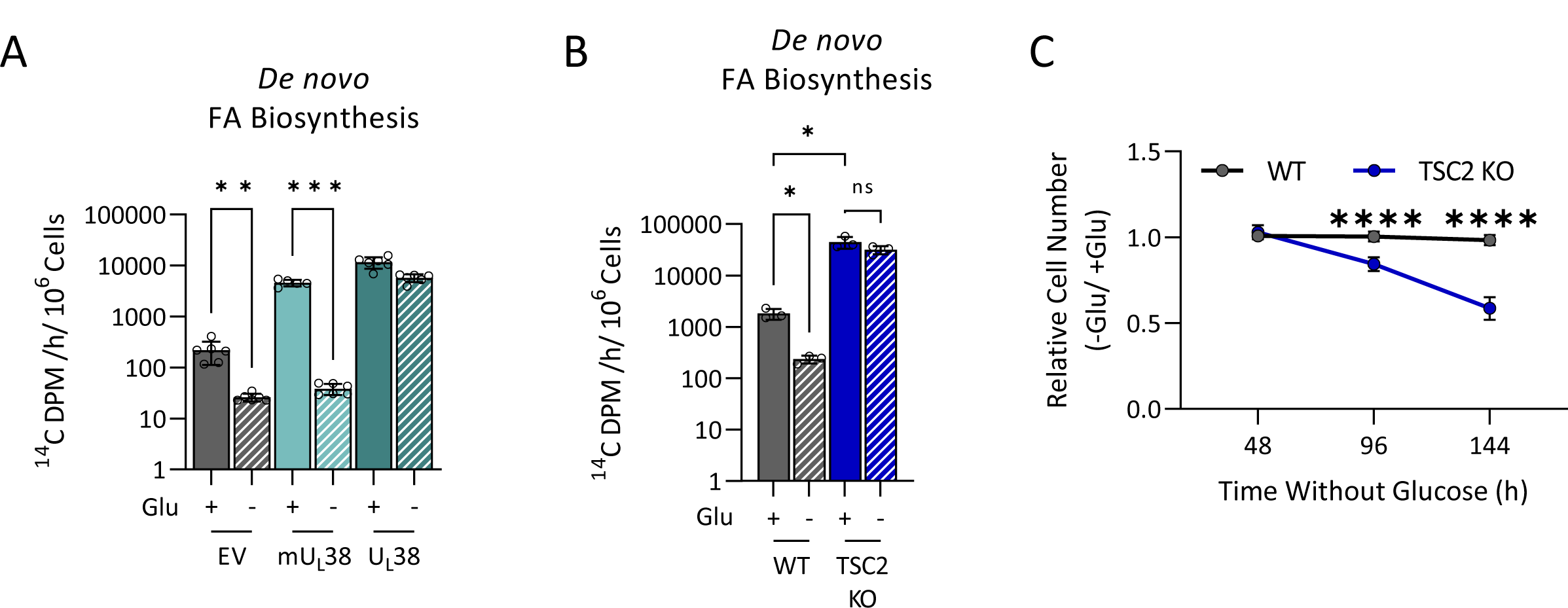
TSC2 inactivation prevents modulation of fatty acid biosynthesis upon glucose limitation. **(A)** MRC-5 fibroblasts expressing an empty vector (EV), wild-type UL38 (UL38), or mutant (mUL38) were grown to confluence and challenged with glucose free medium for 16 hours at which time cells were labeled with 14C-acetate for 4h. Lipids were subsequently extracted, saponified and the radioactivity in the lipogenic fraction scintillation counted. Values are means ± SD (n=6). **(B)** MRC5 (WT) and TSC2 knockout fibroblasts (TSC2 KO) were grown to confluence and challenged with glucose free medium for 16 hours at which time cells were labeled with 14C-acetate for 4h and processed for fatty acid biosynthesis as above. Values are means ± SD (n=6) **(C)** MRC-5 (WT) and TSC2 knockout fibroblasts (TSC2 KO) grown in 384-well plates were and challenged with glucose free medium. Nuclei were counted at the indicated times. Values are means ± SD (n=120).

## Discussion

HCMV modulates the cellular metabolic environment to facilitate successful infection. U_L_38 is a major viral determinant that facilitates this pro-viral metabolic environment, for example, by activating glycolysis and inducing amino acid catabolism (Rodríguez-Sánchez et al., 2019). What has been less clear, is whether HCMV-induced metabolic reprogramming results in emergent metabolic vulnerabilities. In this study, we found that U_L_38 expression induces fatty acid biosynthesis. Further, U_L_38-mediated activation of fatty acid biosynthesis is unresponsive to changes in glucose availability, resulting in sensitivity to glucose limitation that can be attenuated with fatty acid biosynthetic inhibitors. U_L_38 mediates this rigid activation of fatty acid biosynthesis via inhibition of the TSC2 tumor suppressor, the inactivation of which also causes rigid activation of fatty acid biosynthesis and sensitivity to glucose limitation (Fig 9). Collectively, these results identify a regulatory circuit between glucose availability and fatty acid biosynthesis that is essential for cell survival under glucose-limited conditions and reveals a metabolic vulnerability associated with both HCMV viral infection and the inactivation of normal metabolic regulatory controls.

**Figure 9.**
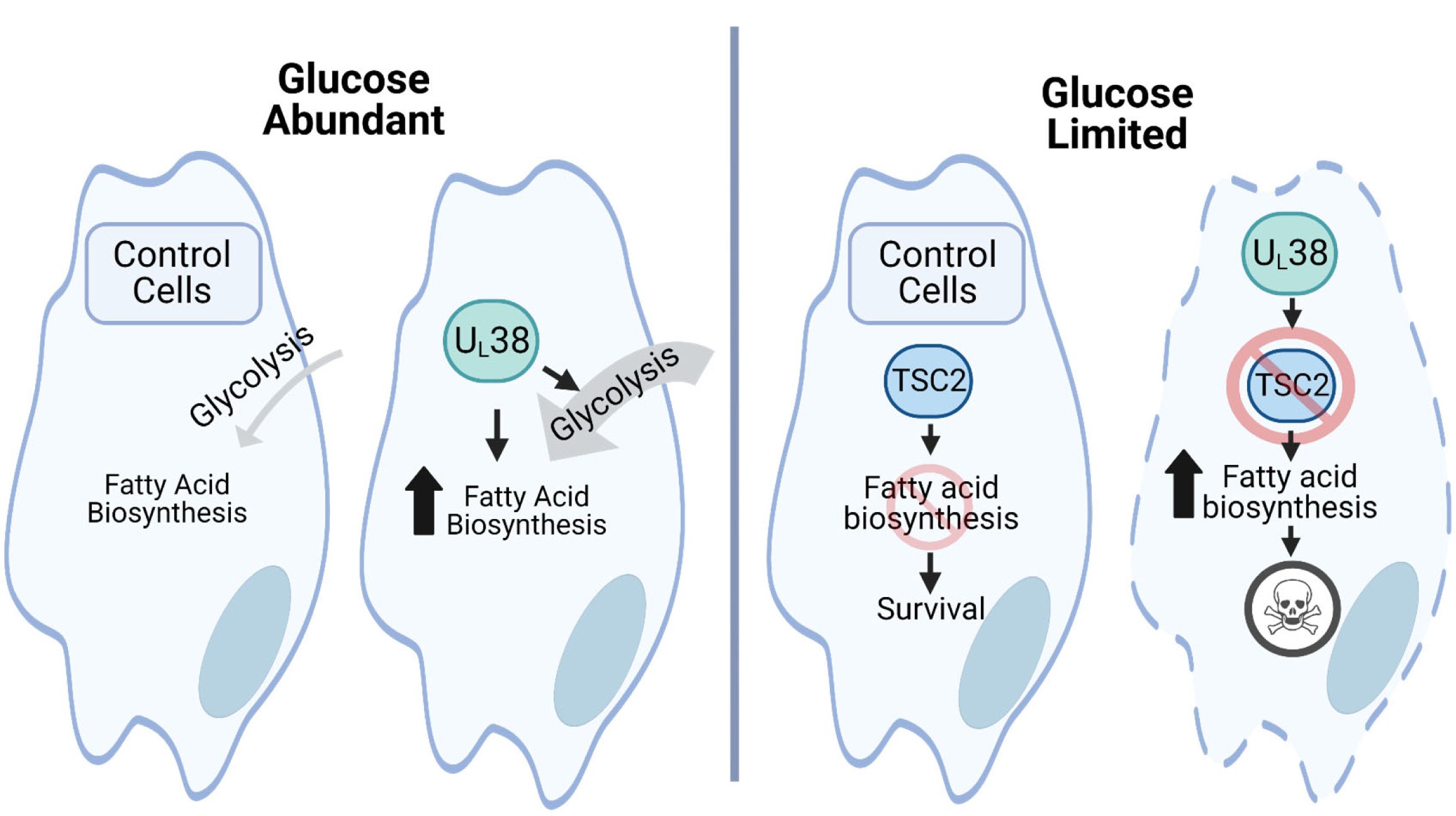
UL38-expression induces rigid fatty acid biosynthetic activation that sensitizes cells to glucose limitation. When glucose is abundant, UL38-expressing fibroblasts have elevated levels of glycolysis and fatty acid biosynthesis. In control cells, when glucose is limited, TSC2 inhibits fatty acid biosynthesis, leading to cell survival. When glucose is limited and UL38 is present, UL38 prevents TSC2 from inhibiting fatty acid biosynthesis, resulting in the loss of membrane integrity and cell death.

We find that U_L_38 significantly increases the expression of numerous metabolism-associated RNA, for example, those involved in glycolysis and fatty acid biosynthesis (Fig 1-2). While it’s unclear how U_L_38 induces these changes, the number of fatty acid and glycolytic genes induced by U_L_38 implicated the potential activation of SREBF1 and HIF1α (Fig 2A). U_L_38 resides primarily in the nucleus at early times during infection (Terhune et al., 2007), and it has been reported to interact with nucleosome remodeling and deacetylation partners (Moorman et al., 2008; Nobre et al., 2019), consistent with a potential role modulating transcription factor activity. However, it remains to be determined whether U_L_38 functions in a transcriptional modulatory role that contributes to the induction of these metabolic target genes.

We find that glycolysis and fatty acid biosynthesis are still induced upon infection with the mU_L_38 HCMV mutant, which possesses T23A/Q24A substitution mutations that attenuate U_L_38-mediated inhibition of TSC2 (Bai et al., 2015; Rodríguez-Sánchez et al., 2019) (Fig 3). Previously, we observed that the expression of mU_L_38 in isolation failed to induce glycolysis, in contrast to the expression of wildtype U_L_38 (Rodríguez-Sánchez et al., 2019). This difference between the expression of mU_L_38 in isolation versus within the context of HCMV infection, suggests that HCMV has evolved redundant mechanisms of glycolytic activation. Possibilities for HCMV-associated activities that induce glycolytic activation, but are U_L_38-independent, include activation of CaMKK and AMPK signaling, which has been shown to activate glycolysis during HCMV infection (Dunn et al., 2021; McArdle et al., 2012; McArdle et al., 2011).

Consistent with a relatively similar metabolic impact between WT and the mU_L_38 HCMV mutant, there was little difference between these two viruses with respect to their replication in fibroblasts (Fig 4E). This likely reflects in part the metabolic remodeling redundancies noted above. In addition, CRISPR-mediated targeting of TSC2 did not rescue the replication of the U_L_38-deletion strain. This is probably indicative of the pleiotropic contributions that U_L_38 makes to infection, including activating mTOR, inducing many facets of central carbon metabolism, blocking ER stress, and inhibiting apoptosis (Moorman et al., 2008; Qian et al., 2011; Rodríguez-Sánchez et al., 2019; Terhune et al., 2007; Xuan et al., 2009). Deletion of the U_L_38 gene severely attenuates HCMV replication *in vitro* (Terhune et al., 2007), and given its pleiotropic functions, it is perhaps unlikely that rescuing any single one of these phenotypes would rescue infection.

Our results indicate that either expression of U_L_38 or the inactivation of TSC2 induces fatty acid biosynthetic activity that is unresponsive to the levels of available glucose. Further, the fatty acid biosynthetic rigidity associated with U_L_38 was reversed with a mutant U_L_38 with reduced TSC2 interaction. These findings suggest that TSC2 plays an important role in modulating fatty acid biosynthetic activity in response to glucose levels. This could reflect an important role of TSC2 for mTOR inhibition in the absence of glucose. TSC2 is a well-known inhibitor of mTOR (Inoki et al., 2003), which can induce fatty acid biosynthesis via the activation of SREBP transcription factors (Eberle et al., 2004; Porstmann et al., 2008). Alternatively, TSC2 could be acting independently of mTOR as mTOR-independent activities have been described (Alves et al., 2015; Brugarolas et al., 2003; Gan et al., 2008; Rodríguez-Sánchez et al., 2019). However, to date, mTOR-independent TSC2 links to fatty acid biosynthesis are not well described. Additionally, it remains to be determined whether the decoupling of glucose availability and fatty acid biosynthesis represents a single metabolic circuit broken by TSC2 inactivation, or whether it may reflect a broader anabolic rigidity or insensitivity to a variety of nutrient cues, for example, the availability of important amino acids such as glutamine or methionine.

Sensitivity to glucose limitation, i.e., glucose addiction, is a common vulnerability associated with cancerous cells (Jiang et al., 2011; Warburg, 1956). It remains to be determined whether the linkage we observe between glucose sensitivity and fatty acid biosynthetic rigidity is maintained more broadly, for example in cancerous cells. Notably, mTOR activity is activated by mutations in a variety of oncogenes and tumor suppressors, thus suggesting the possibility that TSC2 might be a commonly mutated tumor suppressor. On the contrary, TSC2 mutations are not very common, with TSC2 loss reported in 0.1% of all cancers and TSC2 point mutations present in 3.39% of tumors (“AACR Project GENIE: Powering Precision Medicine through an International Consortium,” 2017). The relative lack of cancer-associated inactivating TSC2 mutations may result from the increased nutrient sensitivity associated with TSC2 loss and the inability of cells with TSC2 mutations to reconcile the metabolic needs for anabolic growth with nutrient availability (Choo et al., 2010).

Cells must regulate a variety of anabolic activities based on the availability of various nutrients. Viral infection and oncogenesis remodel cellular metabolic controls in ways that sensitize cells to metabolic perturbation, raising the possibility that these vulnerabilities can be targeted for therapeutic intervention. Increasing our understanding of the molecular mechanisms through which nutrient concentrations are sensed and subsequently how this information is conveyed to regulate anabolic enzymes will likely provide novel strategies to exploit these metabolic vulnerabilities for therapeutic purposes.

## Materials and Methods

### Cell culture, viruses, and viral infections

Human telomerase immortalized fibroblasts (MRC5-hT) and HEK 293T cells were cultured in Dulbecco’s Modified Eagle Media (DMEM; Gibco) supplemented with 10% (vol/vol) fetal bovine serum (FBS; Atlanta Biologicals) and 1% penicillin-streptomycin (Pen-Strep; Life Technologies) at 37 °C in a 5% (vol/vol) CO_2_ atmosphere. For glucose limitation assays, fibroblasts were cultured in glucose-free DMEM (Gibco) supplemented with 1% Pen-Strep. UL38 and mUL38 expressing MRC5 were generated via lentiviral transduction as indicated below. Unless indicated otherwise, the wild-type viral infections were performed using a virus derived from a BADwt bacterial artificial chromosome (BAC) clone of the HCMV AD169 laboratory strain. The recombinant HCMV-ΔU_L_38 BAC derived virus which lacks the entire U_L_38 allele was generously provided by Thomas Shenk, Princeton University (ΔU_L_38). AD169 with mutant U_L_38 allele (mU_L_38) was generated using BAC recombineering as previously described using primers for KanR insertion and gBlocks® Gene Fragment (IDT) containing the U_L_38 T23A/Q24A mutation. Viral stocks of AD169 WT and AD169 mUL38 were propagated in MRC5-hT while AD169 ΔU_L_38 were propagated in U_L_38-expressing MRC5-hT. The titers of viral stocks were determined by plaque assay analysis in MRC5-hT cells. For viral infection assays, cells were grown to confluence upon which medium was removed, and serum-free medium was added. Cells were maintained in a serum-free medium for 24 h before infection at which point they were either mock infected or infected at a multiplicity of infection of 3.0 pfu/cell. After a 2 h adsorption period, the inoculum was aspirated, and fresh serum-free medium was added. For the assessment of glucose-induced vulnerabilities, cells were synchronized with serum-free media for 24h, prior to media change to glucose-free DMEM. For analysis of viral titers, infectious virus was quantified via serial dilution and IE1 expression analysis in MRC5-hT cells at 48 hpi unless otherwise indicated.

### Lentiviral transduction and cell line generation

Lentiviral vectors for U_L_38 and mU_L_38 were generated via cloning with Gibson assembly. Wild type TB40/e U_L_38 allele (U_L_38) was amplified by PCR from the TB40/e BAC clone (EF999921.1) using the following primers: 5’-CTTTAAAGGAACCAATTCAGTCGACTGGATCATGACTACGACCACGCATAGCACCGCCGC-3’, 5’-AACCACTTTGTACAAGAAAGCTGGGTCTAGCTAGACCACGACCACCATCTGTACCACGTC-3’.

A TB40/e mutant U_L_38 allele-T23A/Q24A (mU_L_38) was synthesized as a 996bp gBlocks® Gene Fragment (IDT) using the TB40/e U_L_38 sequence (EF999921.1) and mutating the sequence corresponding to the 23 and 24 amino acids. This construct was amplified by PCR using the same primers described above for the wild-type U_L_38 allele. Both U_L_38 and mU_L_38 constructs were then cloned via Gibson assembly into a pLenti CMV/TO Puro plasmid (Addgene plasmid 22262) that was digested with BamHI and XbaI. pLenti CMV/TO/Puro/empty (EV) was provided by Hartmut Land, University of Rochester. The resulting vectors were then used for lentiviral transduction. Pseudo-type lentiviruses were produced in 10 cm dishes containing 293T cells (∼50% confluent) and transfected with 2.6 μg lentiviral vector, 2.4 μg PAX2, and 0.25 μg vesicular stomatitis virus G glycoprotein expression plasmids using the Fugene 6 transfection reagent (Promega) according to the manufacturer’s recommendations. Twenty-four hours after transfection, the medium was replaced with 4 mL of fresh medium. Lentivirus– containing medium was collected after an additional 24 h and filtered through a 0.45 μm filter and applied to MRC5-hT fibroblast in the presence of 5 μg/mL polybrene (Millipore) and incubated overnight. The lentivirus-containing medium was then removed and cells were allowed to recover for 72 hours whereupon cells were passaged in the presence of 1 μg/mL puromycin (VWR) and remained under selection for 3 days.

TSC2 knockout and control cell lines were generated in MRC5-hT fibroblasts using a CRISPR-Cas9 ribonucleoprotein (RNP) system. Guide crRNA targeting TSC2 (ACAAUCGCAUCCGGAUGAUA, CACAAAUCUGCCCUAUCAUC, GUGGCCUCAACAAUCGCAUC) or a RelA (GAUCUCCACAUAGGGGCCAG, positive control) were obtained from IDT. To make functional guide RNA, crRNA was combined with tracrRNA (IDT). The ribonucleoprotein complexes were then prepared by combing guide RNA and Cas9 2NLS protein (Synthego) using a 9-to-1 guide-to-protein ratio. The RNPs were then electroporated into fibroblasts using the Neon electroporation system (Thermo) following the manufacturer’s protocol and using the following electroporation settings: 1 pulse, 1100 volts, 30 ms pulse width. Cas9 alone without a guide was also electroporated into cells as a negative control. After electroporation cells were allowed to recover and then clonal selection was performed. Clonal populations were expanded and then knockdown efficiency was determined by protein and DNA analysis. Western blotting for each clone was done to identify clones without detectable protein levels prior to DNA sequencing. Genomic DNA was amplified using primers specific to the edited region of TSC2 (AAAGTGCAGGGATTACAGGCCT, TGGAAATGGGGAGCAAAGGGAT; IDT). The amplicon was purified and the sequence was determined by Sanger sequencing which was performed by Genewiz. Synthego ICE analysis was used to deconvolute sequencing reads to determine KO efficiency. (Fig. S2-S3). For experiments, TSC2 KO cell-line 1, was only used for HCMV viral growth curves. TSC2 KO cell line 2 was used for the viral growth curve and all other TSC2 KO experiments.

### RNA sequencing and analysis

For the RNA sequencing experiments, EV or U_L_38 expressing cells were grown to confluence in 10 cm dishes and synchronized with serum-free media for 24 hrs. Total cellular RNA was extracted using TRIzol reagent (Invitrogen) following the manufacturer’s instructions. DNase was used to treat each sample and remove any residual DNA whereupon the RNA was purified using a RNeasy MinElute cleanup kit (Qiagen). Subsequently, the RNA concentration and quality were determined using a NanoDrop 1000 spectrophotometer and Agilent Bioanalyzer, respectively. Library construction and next-generation sequencing were performed by the University of Rochester Genomics Research Center (GRC). Libraries were constructed using the TruSeq stranded mRNA sample preparation kit (Illumina). Briefly, oligo(dT) magnetic beads were utilized to purify mRNA from 200 ng total RNA. Random hexamer priming was used for first-strand cDNA synthesis, followed by second-strand cDNA synthesis using dUTP incorporation for strand marking. Double-stranded cDNA then underwent end repair and 3′ adenylation. cDNA was modified by the ligation of Illumina adapters to both ends. Samples were purified by gel electrophoresis and PCR primers specific to the adapter sequences were used to generate cDNA amplicons ranging in size from 200 to 500 bp. The University of Rochester Genomics Research Center (GRC) prepared the library and performed the sequencing as previously described (Goodwin et al., 2019). DESeq2 was used to perform data normalization and differential expression analysis with a false discovery rate (FDR) adjusted p-value threshold of 0.05 on each set of raw expression measures. For gene set enrichment analysis, genes with a log_2_ expression greater than 0.6 and FDR-adjusted p-value of less than 0.05 were used. Pathway enrichment was performed using the PANTHER classification system (Mi et al., 2019) and transcription factor enrichment was performed using the TRRUST reference database (Han et al., 2015). All enrichment analysis was conducted utilizing the Enrichr website from the Mayan Lab (Chen et al., 2013; Xie et al., 2021). RNA-seq data were submitted to the NCBI gene expression omnibus repository (GSE232603).

### Real-time qPCR

Total cellular RNA was extracted using TRIzol (Invitrogen) and following the manufacturer’s protocol. The purified RNA was then used to generate cDNA using SuperScript II reverse transcriptase (Invitrogen) and random hexamer priming according to the manufacturer’s instructions. Primers for each gene were obtained from IDT. Transcript abundance was measured by real-time PCR analysis using Fast SYBR green master mix (Applied Biosystems), and QuantStudio5 system (Applied Biosystems). Gene expression equivalent values were determined using the 2^− ΔΔCT^ method. Multiple reported “housekeeping genes” had increased RNA abundance in U_L_38-expressing cells during RNAseq analysis, including GAPDH, so values were normalized to LAMC1 levels. The primers that were used for real-time PCR are listed below:

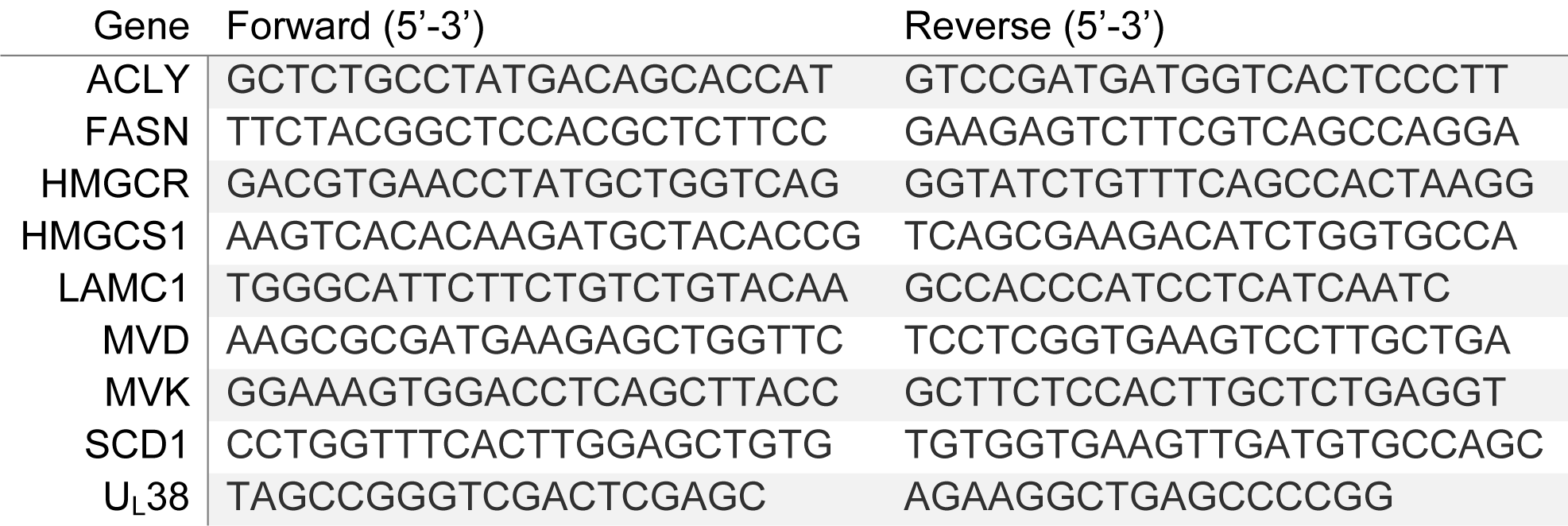

### Fatty acid biosynthesis assays

Measurements of fatty acid biosynthesis were performed by measuring the incorporation of (1-^14^C) acetic acid (Perkin Elmer) into cellular lipids as described previously(Spencer et al., 2011). Fibroblasts were seeded in 12-well plates and grown to confluence at which point the cells were synchronized with serum-free DMEM. Cells were maintained in this serum-free medium for 24 h. At this point cells were infected at an MOI of 3 pfu/cell, given a 1-hour adsorption period, and then provided with fresh serum-free media. Infected cells were then incubated for 44 hours before proceeding. For cells that were not infected, after 24 hours in serum-free media, the media were aspirated, cells were rinsed with DPBS, and fresh serum-free medium was added either with or without glucose and incubated for 20 hours. For both infected and uninfected cells, media were again removed and replaced with media supplemented with 1 μCi of 56.2 mCi mmol^−1^ (1-^14^C) acetic acid. Cells were incubated at 37 °C for 4 hours at which point they were rinsed with PBS and dissolved in 0.5 mL of 0.2 M KOH. Samples were then saponified by the addition of 0.3 mL of 50% KOH and 2.0 mL of ethanol, followed by a 2-h incubation at 75°C and then an overnight incubation at room temperature. After saponification, the samples were extracted two times with 4.5 mL of hexane. The hexane-containing fractions were discarded, and the remaining fractions containing fatty acids were acidified by the addition of 0.5 mL 12 M HCl. After acidification, the samples were extracted in 4.5 mL hexane. The organic fractions were dried under N_2_, resuspended in 50 μL of 1:1 chloroform-hexane, combined with 5 mL of Ecoscint O liquid scintillation fluid, and assessed for radioactivity using a Beckman LS6500 liquid scintillation counter.

### Cell viability quantification

Crystal violet was used to assay cell viability during HCMV infection. MRC5-hT cells were grown to confluence, resulting in ∼3.2 × 10^4^ cells per cm^2^. Once confluent, the medium was removed, and serum-free medium was added. Cells were maintained in a serum-free medium for 24 hours before infection at which point they were mock infected or infected at a multiplicity of infection of 3.0 pfu/cell. After a 2-hour adsorption period, the inoculum was aspirated, and new serum-free media containing or free of glucose were added to the plates. Cell viability was monitored by eye every 24 h and imaged as the infections continued. After 72 h media were aspirated and cells were fixed in 10% neutral buffered formalin (NBF) (VWR) for 20 minutes (min). Fixed cells were washed one time with water and stained with 0.1% crystal violet for 20 min, after which the cells were washed three times with water and allowed to dry. Crystal violet dye was then solubilized by adding 2 mL of 10% acetic acid to each well and incubated for 20 min with shaking. Samples were diluted 1:4 in water and quantified by the absorbance at 595 nm.

### LDH Release Assay

As a measure of cellular membrane integrity, an LDH release assay was performed following the manufacturer’s protocol (Promega). Briefly, EV and U_L_38 expressing cells were seeded in 96-well plates and grown to confluence. Cells were synchronized with serum-free media and incubated for 24 hours before changing the media to media either containing or free of glucose. For LDH release assays with fatty acid biosynthesis inhibitors, the compounds were added with the media change. Cells were incubated at 37 °C for up to 72 hours. At 12, 24, 48, and 72 hours post media change, 10 μL aliquots of media were removed and diluted 100-fold in LDH storage buffer (200mM Tris-HCl (pH 7.3), 10% Glycerol, 1% BSA). The dilutions were added to a 96-well plate and combined with LDH assay buffer, incubated for 30 minutes at room temperature, and luminescence was measured using a Tecan Spark microplate reader.

### Compound library and high-throughput compound screening

The compound library for the high-throughput screen was prepared as described previously (Harris et al., 2019). Before screening with the compound library, a Z’ factor was measured to determine the robustness of our assay. (Zhang et al., 1999). Each well of a 384-well pate (Corning) was seeded with 6,000 cells in a volume of 50 µL using a Multidrop Combi reagent dispenser (Thermo Scientific). Cells were incubated at 37 °C for 24 hours at which point they were synchronized with serum-free DMEM and incubated for an additional 24 hours. Cells then underwent a media exchange in which half the plates received the normal DMEM while the other half received glucose-free DMEM (Gibco). To perform the media exchange, media were aspirated down to 25 µL and then 75 µL of target medium was added. This was repeated 6 times. After the final time, media were aspirated down to 25 µL, and 25 µL of media were added bringing the well volume to 50 µL. Cells were then incubated at 37 °C for 96 hours whereupon they were fixed with 4% paraformaldehyde and nuclei stained with Hoescht. Cells were then quantified as described below and the average and standard deviation of the number of cells were used to determine the Z’ factor of 0.49. To determine compounds that inhibit the sensitivity to glucose withdrawal, cells were seeded, synchronized and glucose deprived as described above. After switching media to glucose-free media, 100 nL of compounds from the library plates were pin transferred onto the cells and incubated at 37 °C for 96 hours, at which point cell numbers were quantified as outlined below. Data post-processing was conducted using an R script, followed by analysis in GraphPad Prism.

### Measurement of metabolite levels

For the quantification of steady-state metabolite levels, five 10-cm dishes were seeded with either U_L_38 or EV MRC5-hT. Once confluent, the medium was replaced with serum-free medium. Cells were maintained in this serum-free medium for 24 h, at which time the media were aspirated, cells were rinsed with DPBS, and fresh serum-free media was added either with or without glucose. After an additional 24 hours cells were harvested for LC-MS/MS analysis, protein abundance, or cell counts. For LC-MS/MS analysis, metabolic activity was quenched by adding dry ice temperature 80:20 OmniSolv® methanol:water. Cells were incubated at −80°C for 10 min and then scraped while being maintained on dry ice. All samples were vortexed thoroughly and centrifuged at full speed for 5 minutes to pellet insoluble materials. The supernatants were saved and the pellets were reextracted twice more with 500 uL 80:20 methanol:water. After pooling all three extractions, 500 µL were removed and the remaining 4.5 mL were completely dried under N_2_ gas. After drying, samples were resuspended in 200 µl of 80% methanol. For amino acid quantification, 100 µl of the above methanol dilutions were derivatized with 1 µl benzyl chloroformate and 5 µl triethylamine. The samples were then centrifuged at 4°C for 5 minutes at full speed to pellet insoluble material and subsequently analyzed by LC-MS/MS as indicated below.

### Quantification of cell numbers

For cell culture experiments in 96-well (Greiner Bio-One, #655160) or 384-well (Corning, #3764) plate formats, cells were washed with PBS (Thermo Fisher, BP399-20), fixed, and stained using a solution of 5 µg/mL Hoechst 33342 (Thermo Fisher) and 4% formaldehyde (Sigma). Cells were then imaged using a Cytation 5 automated imaging reader (BioTek). Each well was imaged using a 4X magnification objective lens and predefined DAPI channel with an excitation wavelength of 377 nm and emission wavelength of 447 nm. Gen5 software (BioTek) was used to determine cell number by gating for objects in the DAPI channel with a minimum intensity above a background of 3000 and a size between 10 and 75 µm.

### LC-MS/MS Analysis and Normalization

Metabolites were analyzed using reverse phase chromatography with an ion-paring reagent in a Shimadzu HPLC coupled to a triple quadrupole mass spectrometer running in negative mode with selected-reaction monitoring (SRM) specific scans as previously described (DeVito et al., 2014; Smith et al., 2016). Briefly, LC-MS/MS was performed by using a LC-20AD HPLC system (Shimadzu) and a Synergi 4 μm Hydro-RP 80A 50×2mm column(Phenomenex), and the LC parameters were as follows: autosampler temperature, 4 °C; injection volume, 10 μL; column temperature, 40 °C; and flow rate, 0.5 mL/min. The LC solvents were: solvent A, 100% methanol; and solvent B, 10 mM tributylamine and 15 mM acetic acid in 97:3 (vol:vol) water:methanol. The gradient conditions were as follows (vol/vol): negative mode-t = 0, 85% B; t = 4, 3%B; t = 5, 3% B; t = 5.1, 85% B. Mass-spectrometric analyses were performed on a TSQ Quantum Ultra triple-quadrupole mass spectrometer running in multiple reaction monitoring mode (Thermo Fisher Scientific). LC-MS/MS data were analyzed using the publicly available mzRock machine learning toolkit (http://code.google.com/p/mzrock/), which automates SRM/HPLC feature detection, grouping, signal-to-noise classification, and comparison to known metabolite retention times (Melamud et al., 2010).

For the relative quantification of intracellular metabolite levels, protein-normalized peak heights were normalized by the maximum value for a specific metabolite measured across the samples run on a given day. This normalization serves to reduce the impact of inter-day mass spectrometry variability, i.e. batch effects while preserving relative differences between samples. Protein-normalized metabolite concentration data were utilized for principal component analysis performed using the publicly available software MetaboAnalyst 3.0 (http://www.metaboanalyst.ca) (Xia et al., 2015).

### Protein analysis and western Blotting

For protein analysis, cells were harvested from fibroblasts using RIPA buffer supplemented with Pierce Protease Inhibitor tablets (Thermo Scientific). Cell extracts were then sonicated and either used for protein quantification via Bradford assay or used for blotting. For western blotting, samples were treated with Laemmli SDS sample buffer (4X; Bio-Rad) with 5% β-mercaptoethanol (Sigma) and then heated for 5 min at 100 °C. Debris was pelleted via centrifugation at 14,500 RPM for 5 minutes, run on 4-20% polyacrylamide gels (Genescript), and transferred onto 0.2 µm nitrocellulose membranes (Bio-Rad). Membranes were then stained with Ponceau S to ensure equivalent protein loading and transfer. Blots were then blocked by incubation in 5% milk in TBST (50 mM Tris-HCl, pH 7.6, 150 mM NaCl, 0.1% Tween 20) for 1 hour and subsequently reacted overnight with indicated antibodies in TBST with 0.05% bovine serum albumin (BSA). Excess primary antibodies were rinsed away with TBST and reacted with secondary antibodies in TBST and 0.05% BSA for 1 hour before being rinsed again in TBST. The signal was visualized using an enhanced chemiluminescence (ECL) system (Bio-Rad) and imaged using the Molecular Imager Gel Doc XR+ system (Bio-Rad). The following antibodies were used for immunoblot analysis: Cleaved Caspase 3 (Cell Signaling, 9661), Vinculin (Cell Signaling, 13901), tuberin (TSC2; Santa Cruz Biotechnology, glyceraldehyde-3-phosphate dehydrogenase (GAPDH; Cell Signaling, 2118), anti-U_L_38 (8D6), anti-IE1(Zhu et al., 1995).

### Statistical Analysis

For mass spectrometry data analysis, protein-normalized peak heights were analyzed using the publicly available software MetaboAnalyst 3.0 (http://www.metaboanalyst.ca). The data were auto-scaled, i.e., mean-centered and divided by the standard deviation of each variable, followed by Principal Component Analysis (PCA) multivariate analysis and hierarchical clustering. Statistical analyses were completed using GraphPad Prism 9.0. Statistical significance was determined by one-way ANOVA. Post hoc analysis was performed using the Fisher’s least significant difference (LSD) method.

## Acknowledgments

We would like to thank John Ashton and Tim Bushnell for their support regarding the frequent use of institutional high-throughput screening equipment. This research has also been facilitated by the services provided by the University of Rochester Genomics Research Center (GRC). The work was supported by NIH grants (AI127370 and AI50698) awarded to J.M., the American Association for Cancer Research and Breast Cancer Research Foundation (20-20-26-HARR), and the Breast Cancer Coalition of Rochester to I.S.H. M.H.R. was supported by the T32 training grant in Cellular, Biochemical and Molecular Sciences (GM068411) and the Elon Huntington Hooker dissertation fellowship. I.R.-S. was supported by a predoctoral fellowship from the American Heart Association.

## Author contributions

M.H.R., I.R.-S., I.S.H., and J.M. designed research; M.H.R., I.R.-S., and X.S. performed research; M.H.R., I.R.-S., X.S., I.S.H., and J.M. analyzed data; and M.R., and J.M. wrote the paper.

## Competing interests

The authors declare that they have no known competing financial interests or personal relationships that could have appeared to influence the work reported in this paper.

